# Muscle oxygenation maintained during repeated sprints despite inspiratory muscle loading

**DOI:** 10.1101/599936

**Authors:** Ramón F. Rodriguez, Nathan E. Townsend, Robert J. Aughey, François Billaut

**Author notes:** These authors contributed equally to this work. Corresponding author, Prof François Billaut, Department of kinesiology, Faculty of medicine, University Laval, 2300 rue de la Terrasse, Quebec (QC) G1V 0A6, (FB).

## Abstract

A high work of breathing can compromise limb oxygen delivery during sustained high-intensity exercise. However, it is unclear if the same is true for intermittent sprint exercise. This project examined the addition of an inspiratory load on locomotor muscle tissue reoxygenation during repeated-sprint exercise. Ten healthy males completed three experimental sessions of ten 10 s sprints, separated by 30 s of passive rest on a cycle ergometer. The first two sessions were “all-out’ efforts performed without (CTRL) or with inspiratory loading (INSP) in a randomised and counterbalanced order. The third experimental session (MATCH) consisted of ten 10 s work-matched intervals. Tissue saturation index (TSI) and deoxy-haemoglobin (HHb) of the vastus lateralis and sixth intercostal space was monitored with near-infrared spectroscopy. Vastus lateralis reoxygenation (ΔReoxy) was calculated as the difference from peak HHb (sprint) to nadir HHb (recovery). Total mechanical work completed was similar between INSP and CTRL (effect size: −0.18, 90% confidence limit ±0.43), and differences in vastus lateralis TSI during the sprint (−0.01, ±0.33) and recovery (−0.08, ±0.50) phases were unclear. There was also no meaningful difference in ΔReoxy (0.21, ±0.37). Intercostal HHb was higher in the INSP session compared to CTRL (0.42, ±0.34), whilst the difference was unclear for TSI (−0.01, ±0.33). During MATCH exercise, differences in vastus lateralis TSI were unclear compared to INSP for both sprint (0.10, ±0.30) and recovery (−0.09, ±0.48) phases, and there was no meaningful difference in ΔReoxy (−0.25, ±0.55). Intercostal TSI was higher during MATCH compared to INSP (0.95, ±0.53), whereas HHb was lower (−1.09, ±0.33). The lack of difference in ΔReoxy between INSP and CTRL suggests that for intermittent sprint exercise, the metabolic O_2_ demands of both the respiratory and locomotor muscles can be met. Additionally, the similarity of the MATCH suggests that ΔReoxy was maximal in all exercise conditions.

## Introduction

Repeated-sprint exercise is characterised by brief periods of “maximal” exertion, interspersed with incomplete recovery periods. Over the course of a repeated-sprint series, there is a progressive reduction in both peak and mean power output, with a plateau in the latter sprints (1–4). While phosphocreatine (PCr) hydrolysis and anaerobic glycolysis are heavily relied on as a rapid source of adenosine triphosphate (ATP) replenishment in sprint exercise (2, 5), the aerobic system plays an increasingly significant role in maintaining performance when sprints are repeated. In fact, PCr resynthesis and removal of inorganic phosphate are exclusively performed through oxidative processes (6), and sensitive to oxygen (O_2_) availability (7). Maintaining O_2_ delivery to the locomotor muscles during repeated-sprint exercise is therefore an important mediating factor of performance.

Near-infrared spectroscopy (NIRS) offers the possibility to explore O_2_ balance (delivery vs. consumption) in skeletal muscle during sprint activity in real time. Deoxy-haemoglobin ([HHb]) and oxy-haemoglobin ([O_2_Hb]) rise and fall, respectively, proportional to an increase in metabolic activity in the underlying tissue. Relative changes in [HHb] have been primarily examined during repeated-sprint exercise, because this variable is considered to be independent of blood volume (8, 9), and is assumed to provide a reliable estimate of muscle O_2_ extraction (9, 10). At sprint onset, there is a rapid increase in vastus lateralis [HHb], which recovers during rest periods (1, 11, 12). Muscle reoxygenation between sprints may also describe the quality of metabolic recovery (13). Improving this variable has positive effects on repeated-sprint performance (14, 15), whereas a slower reoxygenation is associated with performance impairments (11, 13). Though it is currently unclear if the O_2_ cost of exercise hyperpnea has any influence on locomotor muscle oxygenation trends during repeated-sprint exercise.

The respiratory muscles demand ≈10-15% of total pulmonary O_2_ uptake 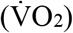 during high-intensity exercise, as well as a considerable portion of cardiac output to maintain adequate O_2_ delivery (16). An elevated work of breathing during high-intensity exercise promotes competition between locomotor and respiratory muscles for available cardiac output (17). In fact, the addition of an inspiratory load to artificially increase the work of breathing during severe exercise (>95% 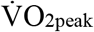) limits endurance capacity via decreased limb perfusion and O_2_ delivery that is mediated by a sympathetically activated vasoconstriction in the locomotor muscles (18, 19). However, at moderate intensities (50-75% 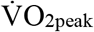) there is no change in vascular resistance or blood flow to the locomotor muscles (20), suggesting that exercise intensity is an important mediator of locomotor vasoconstriction when the work of breathing is high.

It is currently unclear if an elevated work of breathing influences reoxygenation capacity during repeated-sprint exercise. Therefore, we aimed to determine the impact of an elevated work of breathing on 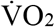, tissue oxygenation trends and mechanical output during repeated-sprint exercise. We believe that by increasing the work of breathing, that there will be no concurrent increase in 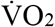, vastus lateralis oxygenation will be compromised and repeat-sprint ability will be impaired.

## Materials and methods

### Subjects

Ten males from a variety of athletic backgrounds (team sports, road cycling, combat sports, CrossFit) were recruited for this study (age 25.5 ± 3.6 years; height 184.00 ± 7.69 cm; body mass 81.45 ± 8.29 Kg; 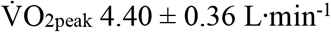, 54.4 ± 5.9 mL·min^-1^·Kg^-1^; 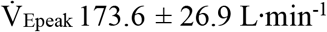). Subject pulmonary function data is presented in Table 1. These subjects were chosen because they were accustomed to producing “all-out” bouts of exercise. Subjects selfreported in a written questionnaire to be healthy non-smokers and with no known neurological, cardiovascular, respiratory diseases, or any other medical conditions. If subjects indicated “yes” to any of the contraindications, they were excluded from participation. After being fully informed of the requirements, benefits, and risks associated with participation, each subject gave written informed consent. Ethical approval for the study was obtained from the institutional Human Research Ethics Committee and the study conformed to the declaration of Helsinki. Prior to each experimental session, subjects were asked to their preceding meal and to refrain from caffeinated beverages for 24 h. Subjects were also asked to refrain from any strenuous exercise for 48 h prior to the experimental sessions.

**Table 1:** Pulmonary function data.

| Measure | Raw | %Predicted |
| --- | --- | --- |
| FVC (L) | $6.0 \pm 0.7$ | $103 \pm 8.8$ |
| FEV<sub>1</sub> (L) | $4.9 \pm 0.5$ | $99.7 \pm 8.1$ |
| FVC/FEV<sub>1</sub> | $79.1 \pm 6.6$ | $95.5 \pm 7.5$ |
| MVV (L·min<sup>-1</sup>) | $212.9 \pm 24.7$ | $110.9 \pm 10.2$ |
| MIP (cmH<sub>2</sub>O) | $-146.6 \pm 19.2$ | |
| MEP (cmH<sub>2</sub>O) | $151.8 \pm 35.8$ | |

Abbreviations are: forced vital capacity, FVC; forced expired volume in 1 s, FEV<sub>1</sub>; maximum voluntary ventilation, MVV; maximum inspiratory pressure, MIP; maximum expiratory pressure, MEP. Values are reported as mean ± SD.

### Experimental design

Subjects visited the laboratory on six occasions. During visit one, subjects completed pulmonary function (spirometry and maximum voluntary ventilation) (Ultima CPX, MGC Diagnostics, Minnesota, USA) and respiratory muscle strength tests (MicroRPM, Micro Medical, Hoechberg, Germany) followed by a maximal ramp exercise test familiarisation. In visit two, a ramp exercise test to volitional exhaustion was completed. During visit three, subjects completed a familiarisation session consisting of the same repeated-sprint protocol used in the experimental sessions. In visits four and five subjects completed the actual repeated-sprint test in a randomised, counterbalanced, cross-over design with either no restriction to their breathing (CTRL), or with inspiratory loading (INSP). During the sixth lab visit, ten work-matched intervals to that of the INSP experimental session were completed (MATCH). The MATCH condition was included in the experiment design to firstly minimise the physiological disturbances that are typically associated with “maximal” sprint performance (2, 3, 5), and to secondly allow for comparison of the data to an exercise trial (INSP) where the same amount of mechanical work was performed by the subjects. All exercise testing was performed on an electronically-braked cycle ergometer (Excalibur, Lode, Groningen, The Netherlands). Experimental sessions were conducted at the same time of day and separated by 3-7 days.

### Maximal ramp exercise testing

A maximal ramp cycling ergometer test was performed to determine 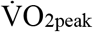. The exercise test was initiated at a work rate of 0 W for 3 min. This was followed by an increase in work rate of 1 W every 2 s (30 W^-1^·min^-1^) until volitional exhaustion or until cadence fell 10 rpm below self selected rate. Expired gases were collected on a breath-by-breath basis (COSMED Quark CPET; Cosmed, Rome, Italy), and peak 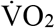 was determined as the highest 30 s average prior to exercise termination. The corresponding 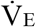 at 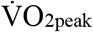 was deemed to be 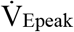. Subjects were familiarised with this protocol during their first visit to the laboratory. Subjects completed the same protocol to volitional exhaustion while wearing a silicone face-mace (Hans Rudolph inc., Kansas, United States of America) that would be used for data collection.

### Repeated-sprint exercise

After arriving at the laboratory, subjects were fitted with NIRS probes and a heart rate monitor. Testing was performed with the cycle ergometer set to isokinetic mode. In this mode, a variable resistance is applied to the flywheel proportional to the torque produced by the subjects to constrain their peddling rate to 120 rpm. Below 120 rpm, no resistance is applied to the flywheel. The handlebars and seat were individually adjusted to each subjects’ characteristics and feet secured using toe cages and retention straps fitted to the ergometer. Crank arm length was standardised to 175 mm. After a 7-min warm-up consisting of 5 min of unloaded cycling at 60-70 rpm and two 4 s sprints (separated by 1 min), subjects rested for another 2.5 min before the repeated-sprint protocol was initiated. The repeated-sprint protocol was ten consecutive 10 s sprints separated by 30 s passive rest (4, 21–23). Subjects were instructed to give an “all-out” effort for every sprint and verbally encouraged throughout to promote a maximal effort. Each sprint was performed in the seated position and initiated with the crank arm of the dominant leg at 45°. Before sprint one, subjects were instructed to accelerate the flywheel to 95 rpm over a 15-s period and assume the ready position 5 s before the commencement of the test. This ensured that each sprint was initiated with the flywheel rotating at ~90 rpm so that subjects could quickly reach 120 rpm. Visual feedback of power output was not available to the subjects during any sprint. The cycle ergometer software provides power and cadence at 4 Hz. Data were exported to Microsoft Excel for analysis. Peak power output was determined by identifying the highest individual power value (watts), which happened to be during Sprint 1 in every case. Mean power output was calculated at the average power within each of the ten sprints. Mechanical work performed (J) was calculated by integrating the power curve over each of the 10 s sprints. Rating of perceived exertion for exercise (RPE_Exercise_), and breathing (RPE_Breath_), was recorded following sprint one, five and ten, using a Borg 6-20 RPE scale. In every case, subjects were asked the questions “how difficult is exercise?” and “how difficult is breathing?”. The scale was anchored so that 6 represented no exertion, and 20 represented maximal exertion.

Inspiratory loading was achieved by placing a plastic disk with a 10-mm opening over the inspiratory side of a two-way non-rebreathing valve (Hans Rudolph inc., Kansas, United States of America) attached to the distal end of the breath-by-breath gas sampling line and turbine (24, 25). The non-rebreathing valve was worn during all the repeated-sprint experimental sessions. The inspiratory load was applied after warm-up, 1 min before the commencement of the repeated-sprint protocol. Work-matched (MATCH) exercise was conducted by performing ten 10 s bouts of exercise separated by 30 s of passive rest and matched for total work performed during the INSP experimental sessions. This was achieved by programming the cycle ergometer for ten bouts of fixed work rate intervals equivalent to each subject’s calculated mean power output for every corresponding sprint repetition (1 to 10) in the INSP experimental sessions. Between each interval intervals during the passive rest period, the cycle ergometer was programmed to 0 watts. The cycle ergometers isokinetic function cannot be used when controlling for power output, therefore, subjects were asked to maintain cadence at 120 rpm during each interval. Warm-up procedures were identical to the CTRL and INSP experimental sessions.

Participants were familiarised with the repeated-sprint protocol during their third visit to the laboratory. Subjects were not instrumented with any devices apart from the silicone face mask and non-rebreathing valve. To expose subjects to the inspiratory loading, they performed the first two sprints of the series with restricted inhalation. The inspiratory loading was promptly removed, and the subjects continued with the remaining eight sprints. Subjects were also asked to rate their perceived exertion for exercise and breathing during this familiarised trial.

### Metabolic and ventilatory measurements

Subjects wore a silicone facemask to which the breath-by-breath gas sampling line and turbine were attached. The analyser was calibrated before each test against known gas concentrations and the turbine volume transducer was calibrated using a 3 L syringe (Cosmed, Rome, Italy). Errant data points due to coughing or swallowing were initially removed and thereafter any breath further than 4 standard deviations from the local mean was removed (26, 27). A 5-breath rolling average was applied for the calculation of peak and nadir for both 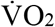 and 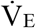 for every 40-s sprint/recovery period to give a single value for each sprint and recovery phase. Inspiratory volume (IV), respiratory frequency (*f*_R_), end-tidal O_2_ partial pressure (P_ET_O_2_), and end-tidal CO_2_ partial pressure (P_ET_CO_2_) were averaged to give one value for each 40-s period. Because the facemask was removed immediately after the tenth sprint, only maximum values were calculated over the first 10 s. Mouth pressure (P_m_) was recorded continuously at 50 Hz with a pressure transducer (Honeywell, New Jersey, United States of America) attached to the saliva port of the non-rebreathing valve via Tygon tubing. Representative data from one subject of the effects of inspiratory muscle loading on P_m_ is displayed in Fig 1. Mean inspiratory and expiratory P_m_ was then calculated as an index of respiratory muscle work. To account for any change in *f*_R_, an index of inspiratory muscle force development was also calculated for each exercise experimental session (∫P_m_ × *f*_R_, expressed in arbitrary units) (28). For statistical analysis, inspiratory P_m_ was converted to positive values and presented in the results as such. Heart rate (HR) (Cosmed, Rome, Italy) and arterial oxygen saturation (estimated by fingertip pulse oximetry; S_P_O_2_) (Nonin Medical, Minnesota, United States of America) was collected on a breath-by-breath basis integrated into the Cosmed system.

**Fig 1.**
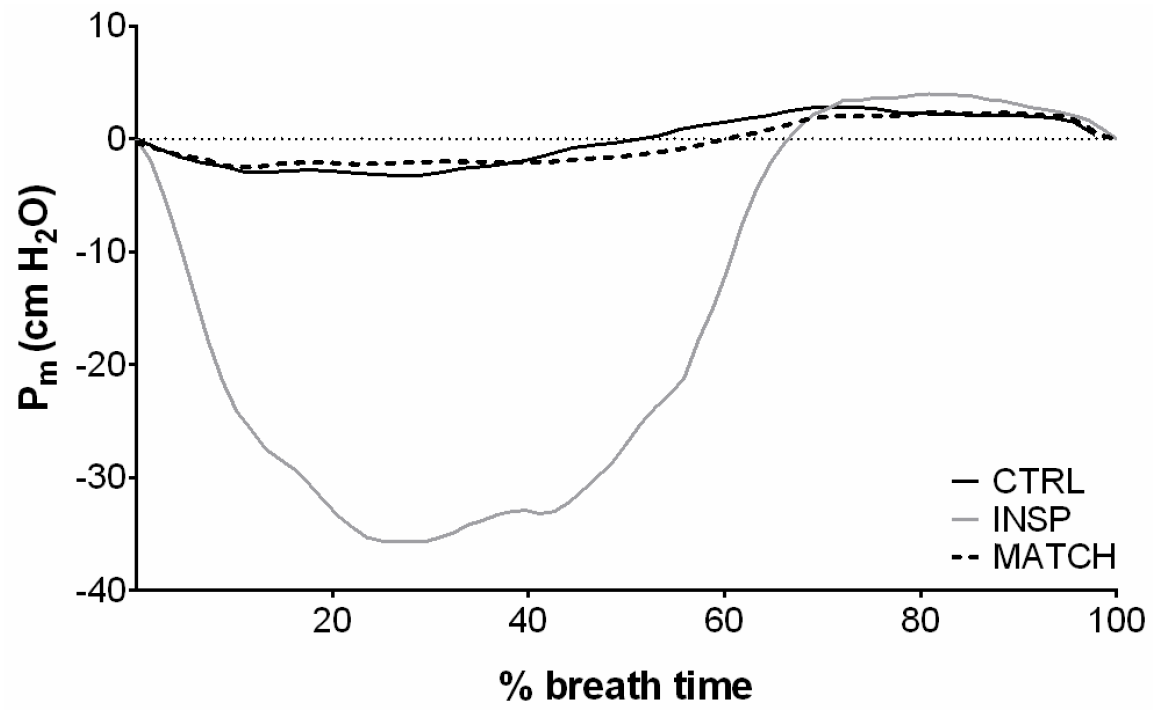
Representative data of the effects of inspiratory muscle loading (INSP) on mouth pressure (Pm) compared to control (CTRL) and work matched (MATCH) exercise conditions.

### Near-infrared spectroscopy

Subjects were instrumented with two NIRS probes to assess muscle oxygenation (Oxymon MKIII, Artinis, the Netherland). The first probe was fixed over the distal part of the vastus lateralis muscle belly approximately 15 cm above the proximal border of the patella of the dominant leg. The second probe was fixed over the left 6^th^ intercostal space at the anterior axillary line of the serratus anterior to assess changes in the accessory respiratory muscles. Probes were held in place with black plastic spacers secured to the skin using double-sided tape and shielded from light using a black self-adhesive elastic bandage. An indelible marker was used to trace the position of the probes to ensure placement can be reproduced in subsequent visits. Optode spacing was set to 4.5 cm and 3.5 cm for vastus lateralis and respiratory muscles, respectively. Skinfold thickness was measured between the emitter and detector using a skinfold calliper (Harpenden Ltd.) to account for skin and adipose tissue thickness covering the muscle. The skinfold thickness for vastus lateralis (1.19 ± 0.69 cm) and respiratory muscles (1.12 ± 0.44 cm) was less than half the distance between the emitter and the detector in every case. A modified form of the Beer-Lambert law was used to calculate micromolar changes in tissue [HHb] and [O_2_Hb] across time using the received optical density from continuous wavelengths of NIR light. A differential pathlength factor of 4.95 was used (12). Tissue saturation index was used as an index of tissue oxygenation (TSI = [O_2_Hb] / ([O_2_Hb] + [HHb]), expressed in %), which reflects the dynamic balance between O_2_ supply and O_2_ consumption in the tissue microcirculation and is independent of near-infrared photon pathlength in tissue. Due to partial data loss in one of the transmitter fibre optic cables for two subjects, respiratory muscle TSI analysis was limited to n = 8. We chose to focus our analysis on Δ[HHb] to allow comparisons to previous research; because Δ[HHb] is independent of changes in total haemoglobin (8); and taken to reflect venous [HHb] which provides an estimate of muscular oxygen extraction (9, 10); and because Δ[O_2_Hb] is influenced by rapid blood volume and perfusion variations caused by forceful muscle contractions (8, 29).

Data were acquired at 10 Hz. A 10^th^ order zero-lag low-pass Butterworth filter was applied to the data to remove artefacts and smooth pedalling induced fluctuations; the resulting output was used for analysis (21). The application of the filter was conducted in the *R* environment (R: A language and environment for statistical computing, Vienna, Austria). Values were then normalised to femoral artery occlusion so that 0% represented a 5-s average immediately prior to the occlusion and 100% represented the maximum 5 s average. Occlusion was performed 3-5 min following the cessation of the sprints while the subjects were supine on an examination bed. Commencement of occlusion was largely influenced by the subject’s wellness following the sprint protocol (e.g. exercise-related syncope, nausea). Subjects were asked to place their foot flat on the examination bed with ≈90° of knee flexion. A pneumatic tourniquet (Rudolf Riester GmbH, Jungingen, Germany) was positioned as high as possible around the thigh and inflated to 350 mm Hg. The tourniquet remained in place until there was a plateau of at least 10 s in vastus lateralis HHb, approximately 5-7 min (9). Tourniquet pressure was monitored continuously to ensure it remained at 350 mm Hg for the duration of the occlusion period. To obtain one value per sprint and recovery for vastus lateralis, peaks and nadirs were identified for each period using a rolling approach (HHb_VL_) (21). Time to peak HHb_VL_ (TTP_HHb_) was also calculated as the time from sprint onset to peak HHb. Reoxygenation capacity (ΔReoxy, %) and reoxygenation rate (Reoxy rate, %·s^-1^) were also calculated as the change from sprint to recovery. The baseline for respiratory muscle HHb was established before warm-up while seated quietly on the cycle. Exercise values are expressed as percent change from baseline. Because there were no clear peaks and nadirs in the respiratory muscle HHb signal, an average was calculated for each 40-s sprint/recovery period (HHb_RM_).

### Statistical analysis

Data in text and figures are presented as mean ± standard deviation. A custom made spreadsheet was used to analyse the effects of INSP and MATCH on laboratory measurements (30). All measures, other than 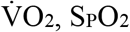, RPE and NIRS responses (except for TTPHHb), were log-transformed before analysis then back-transformed to express the changes in percent units and standardised effects. Relative changes (%) and standardised effects are expressed with 90% confidence limits (90% CL). Practical significance was assessed by calculating Cohen’s d effect size (ES) (31). Standardised effect sizes of <0.2, 0.2-0.5, 0.5-0.8, >0.8 were considered as trivial, small, moderate and large respectively and presented with 90% CL. Probabilities were also calculated to establish if the chance the true (unknown) differences were lower, similar to or higher than the smallest worthwhile change (ES = 0.2). Effects were not considered meaningful if there was <75% probability of being substantially positive/negative relative to the smallest worthwhile change. If the chance of having higher/lower values than the smallest worthwhile difference was both >5%, the true difference was assessed as *unclear.* For clear effects, the likelihood that the true effect was substantial were assessed qualitatively as follows: *likely* (75 to <95%), *very likely* (95-99.5%), *almost certainly* (>99%) (32).

## Results

### Mouth pressure

Mouth pressure responses to exercise and inspiratory muscle loading are presented in Table 2, and representative data from a single subject are presented in Fig 1. Mean inspiratory P_m_ was greater during INSP compared to CTRL (relative difference = 616.6%, 90% CL ±62.2%; ES 13.33, 90% CL ±0.59). Similarly, inspiratory muscle force generation (∫P_m_ × *f*_R_) was *almost certainly* higher during INSP compared to CTRL (753.6% ±91.1%; ES 13.04 ±0.65). But there was an *unclear* difference in mean expiratory P_m_ (−0.6% ±7.1%; ES −0.05 ±0.61).

**Table 2:** Physiological and perceptual responses to repeated-sprint exercise during control (CTRL), inspiratory loading (INSP), and work match (MATCH) conditions.

| Variable | CTRL | INSP | MATCH |
| --- | --- | --- | --- |
| Inspiratory $P_m$ (cm H<sub>2</sub>O) | 2.4 ± 0.3 | 17.6 ± 3.6*** | 1.6 ± 0.3### |
| $\int P_m \times f_R$ (AU) | 76 ± 11 | 658 ± 143*** | 45 ± 9### |
| Expiratory $P_m$ (cm H<sub>2</sub>O) | 2.1 ± 0.2 | 2.1 ± 0.2 | 1.2 ± 0.2### |
| Sprint $\dot{V}O_2$ (% $\dot{V}O_{2peak}$) | 90.7 ± 6.7 | 95.4 ± 4.3 ** | 73.7 ± 6.1 ### |
| Recovery $\dot{V}O_2$ (% $\dot{V}O_{2peak}$) | 70.7 ± 5.2 | 74.9 ± 4.5 * | 58.6 ± 4.3 ### |
| Sprint $\dot{V}_E$ (L·min<sup>-1</sup>) | 161.5 ± 27.5 | 129.1 ± 16.8 *** | 83.4 ± 15.2 ### |
| Recovery $\dot{V}_E$ (L·min<sup>-1</sup>) | 101.4 ± 13.9 | 89.7 ± 11.4 ** | 60.2 ± 10.0 ### |
| IV (L) | 3.0 ± 0.5 | 3.1 ± 0.5 | 2.6 ± 0.4 ### |
| $f_R$ (b·min<sup>-1</sup>) | 48.1 ± 7.8 | 37.8 ± 5.0 *** | 30.2 ± 6.0 ## |
| $P_{ET}O_2$ (mm Hg) | 117.7 ± 2.3 | 113.4 ± 2.8 *** | 105 ± 5.1 ### |
| $P_{ET}CO_2$ (mm Hg) | 35.5 ± 2.3 | 38.6 ± 2.6 *** | 41.6 ± 3.2 ## |
| $SpO_2$ (%) | 97 ± 1 | 97 ± 1 | 97 ± 1 |
| HR (b·min<sup>-1</sup>) | 153 ± 12 | 154 ± 9 | 131 ± 12 ### |
| RPE<sub>Exercise</sub> (AU) |  |  |  |
| Sprint 1 | 15 ± 3 | 14 ± 2 | 12 ± 2 ### |
| Sprint 5 | 17 ± 2 | 17 ± 2 | 13 ± 2 ### |
| Sprint 10 | 18 ± 2 | 18 ± 2 | 13 ± 2 ### |
| RPE<sub>Breath</sub> (AU) |  |  |  |
| Sprint 1 | 12 ± 2 | 15 ± 2 *** | 11 ± 2 ### |
| Sprint 5 | 15 ± 1 | 18 ± 2 *** | 12 ± 2 ### |
| Sprint 10 | 16 ± 2 | 19 ± 1 *** | 12 ± 2 ### |

Abbreviations are: mouth pressure, $P_m$ ; inspiratory muscle force development, $\int P_m \times f_R$ ; pulmonary oxygen uptake, $\dot{V}O_2$ ; pulmonary ventilation, $\dot{V}_E$ ; inspiratory volume, $\dot{V}_E$ ; inspiratory volume, IV; respiratory frequency, $f_R$ ; end-tidal oxygen partial pressure, $P_{ET}O_2$ ; end-tidal carbon dioxide partial pressure, $P_{ET}CO_2$ ; arterial oxygen saturation estimated by pulse oximetry, $S_pO_2$ ; heart rate, HR; rating of perceived exertion for exercise, $RPE_{Exercise}$ ; rating of perceived exertion for breathing, $RPE_{Breath}$ . The symbols represent comparisons between INSP and CTRL (\*), INSP and MATCH (#). The number of symbols; one, two and three denote likely, very likely and almost certainly, respectively, that the chance of the true effect exceeds a small (-0.2-0.2) effect size. Values are reported as mean ± SD.

Mean inspiratory P_m_ was lower during MATCH compared to INSP with a large effect (−91.0% ±0.9; ES −10.18 ±0.42). Likewise, ∫P_m_ × *f*_R_ was lower during the MATCH experimental sessions (−93.1 % ±1.0%; ES −9.92 ±0.52). A large effect was also present for expiratory P_m_ with MATCH being lower than INSP (−40.3% ±5.8%; ES −4.14 ±0.78).

### Mechanical measurements

Total work completed on the cycle ergometer per sprint and over the entire protocol for each condition is presented in Fig 2. There was no meaningful effect of INSP on total work completed over the entire repeated-sprint protocol compared to CTRL (−2.7% ±6.4%; ES −0.17 ±0.42). Similarly, total work performed during MATCH and INSP trials (−0.6% ±0.1%; ES −0.04 ±0.01) were not different. There was no meaningful effect of INSP on peak power output (1097 ± 148 W) compared to CTRL (1158 ± 172 W) (−5.1% ±6.1; ES −0.30, ±0.35). Whereas an *almost certainly* large effect existed between MATCH (773 ± 122 W) and INSP (−29.7%, ±2.3%; ES −2.21 ±0.20).

**Fig 2.**
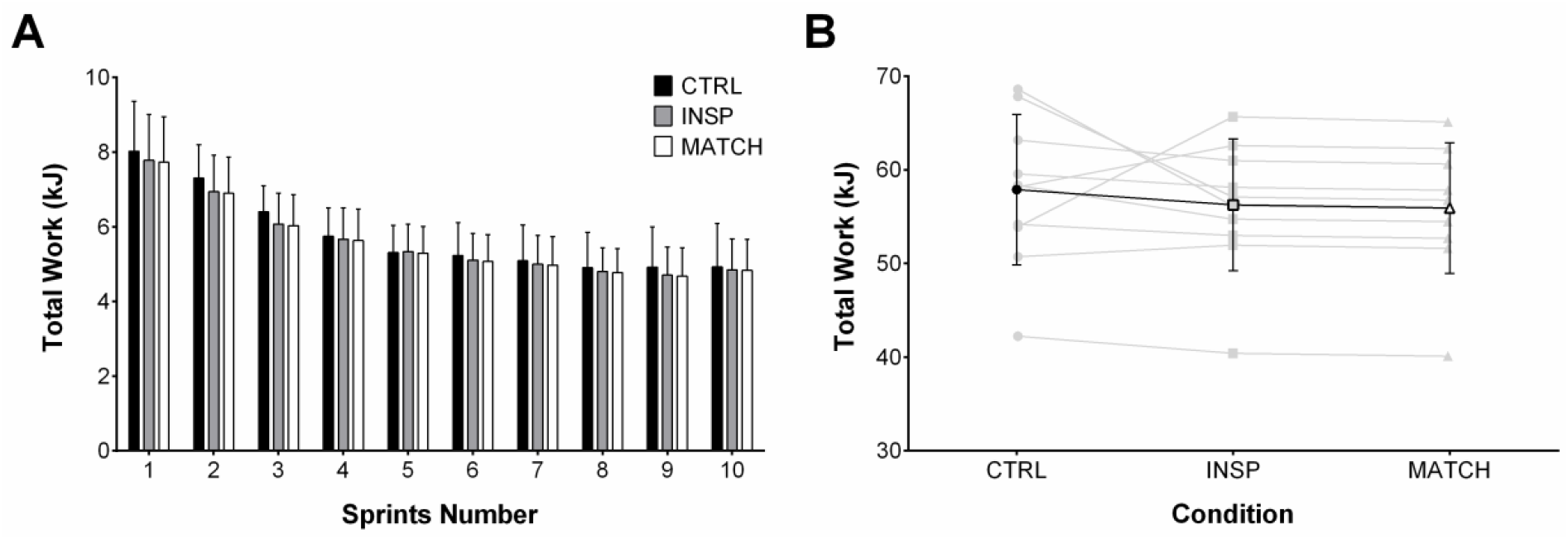
Total mechanical work performed during control (CTRL), inspiratory loading (INSP), and work match (MATCH) exercise conditions. A) Total mechanical work performed during each 10 s sprint. (B) Individual (grey lines) and mean total work (black line) completed over the entire repeated-sprint protocol. Values are presented as mean ± SD.

### Physiological responses

Responses to exercise are presented in Table 2 and Fig 3. Over the entire protocol, 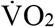 was greater during both sprint (4.7% ±2.7%; ES 0.64 ±0.37) and recovery (4.2% ±3.1; ES 0.74 ±0.55) for INSP compared to CON. Additionally, 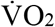 was lower during MATCH compared to INSP during both sprint (−21.7% ±5.0%; ES −4.59 ±1.06) and recovery (16.3% ±2.9%; ES −3.30 ±0.58). Likewise, 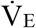 during INSP was lower both during sprint (−19.6% ±3.5%; ES −1.13 ±0.22) and recovery (−11.5% ±5.6; ES −0.80 ±0.41) compared to CTRL. Throughout MATCH, 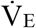 was lower during both sprint (−35.8% ±9.5%; ES −2.92 ±0.98) and recovery (−33.2% ±6.2%; ES −2.81 ±0.65). There was no meaningful difference of IV between INSP and CTRL (2.8% ±4.8%; ES 0.16 ±0.27). On the other hand, IV was *almost certainly* lower during MATCH compared to INSP (−15.6% ±5.1%; ES −0.91 ±0.32). There was an *almost certainly* large effect of INSP on *f*_R_ compared to CTRL (−21.2% ±4.7%; ES −1.41 ±0.35). Additionally, *f*_R_ was *very likely* lower during MATCH compared to INSP (−20.8% ±9.6%; ES −1.53 ±0.79).

**Fig 3.**
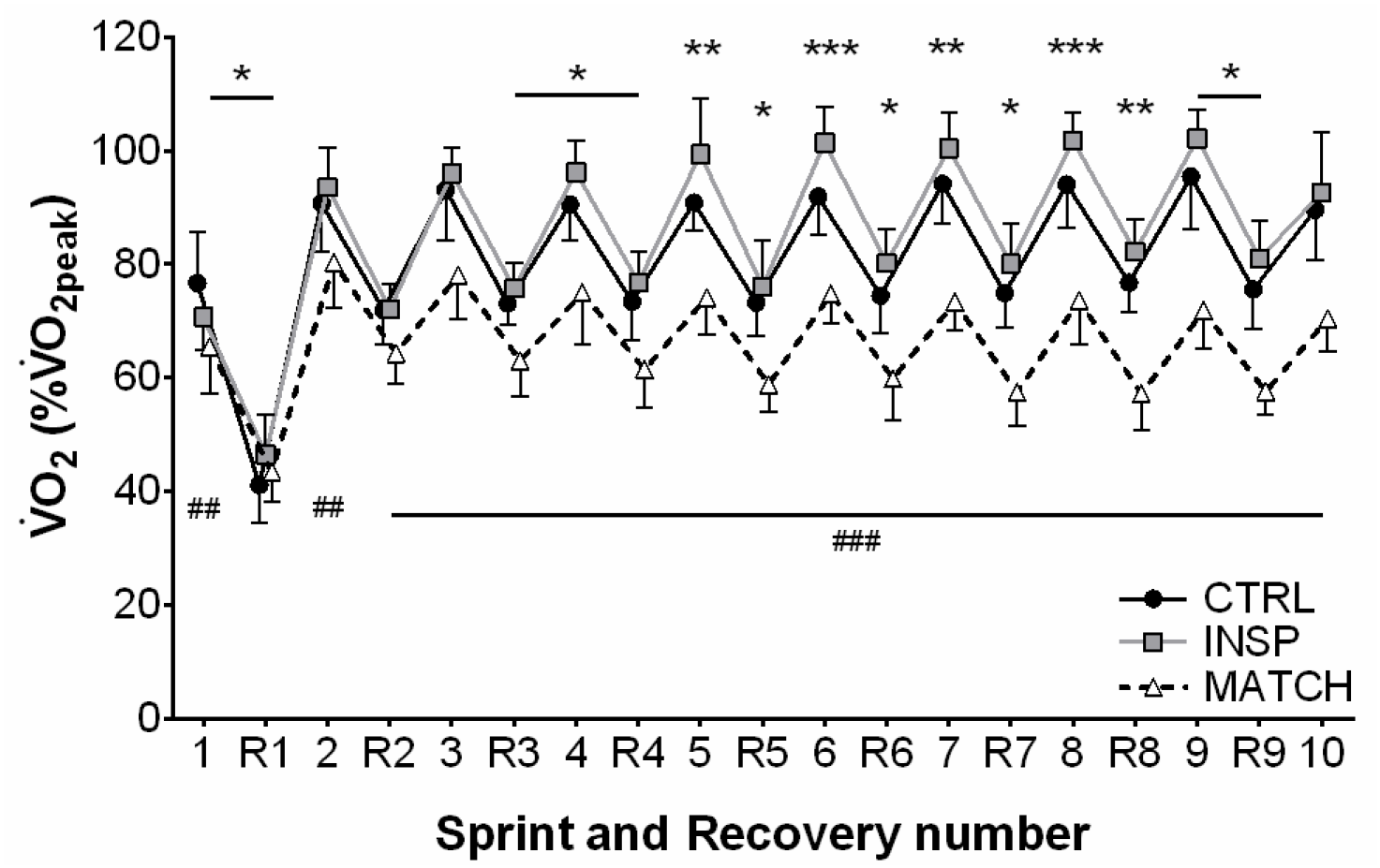
Sprint and recovery pulmonary oxygen uptake 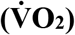 expressed as a percentage of 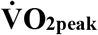 for control (CTRL), inspiratory muscle loading (INSP) and worked matched (MATCH) exercise. The symbols represent comparisons between INSP and CTRL (*), INSP and MATCH (#). The number of symbols; one, two and three denote likely, very likely and almost certainly respectively, that the chance of the true effect exceeds a small (−0.2-0.2) effect size. Vales are presented as mean ± SD.

During INSP, P_ET_O_2_ was lower than CTRL (−3.7% ±2.3%; ES −1.71 ±0.65), and lower during MATCH compared to CTRL (−7.4% ±2.3; ES −2.81 ±0.91). Conversely, P_ET_CO_2_ was higher during INSP compared to CTRL (9.1% ±4.0%; ES 1.22 ±0.51), and higher during MATCH compared to INSP (7.7% ±4.2; ES 1.03 ±0.54). There were *unclear* differences for S_P_O_2_ between both INSP and CTRL (−0.1% ±0.5; ES −0.10 ±0.43), and, MATCH and INSP (0.2% ±0.4%; ES 0.16 ±0.34).

Differences for HR were *unclear* between INSP and CTRL (1.0% ±4.4%; ES 0.11 ±0.50). However, there was a clear *almost certainly* large effect between MATCH and INSP (−15.6% ±3.3; ES −2.66 ±0.61).

Subjects self-report of RPE_Exercise_ during INSP was *likely* trivial after sprint 1 (−2% ±0.5%; ES −0.08 ±0.21) compared to CTRL, and *unclear* after sprint 5 (0.5% ±1.1%; ES 0.14 ±0.52) and sprint 10 (0.4% ±1.0%; ES 0.19 ±0.49). Whereas RPE_Breath_ was *almost certainly* – higher after sprint 1 (2.8% ±0.7%; ES 1.46 ±0.37), sprint 5 (2.4% ±1.0; ES 1.79 ±0.77), and sprint 10 (2.7% ±1.2%; ES 1.32 ±0.57). Comparing MATCH to INSP, RPE_Exercise_ was *almost certainly* lower after sprint 1 (−2.1% ±0.8%; ES −0.98 ±0.37), sprint 5 (−4.4% ±1.3%; ES −2.78 ±0.79), and sprint 10 (−5.6% ±1.2; ES −3.43 ±0.73). Similarly, RPE_Breath_ was *almost certainly* lower after sprint 1 (−4.0% ±1.1%; ES −1.69 ±0.46), sprint 5 (−6.1% ±2.0%; ES −2.36 ±0.78), and sprint 10 (−6.8% ±1.6%; ES −3.93 ±0.95)

### Muscle oxygenation

Muscle oxygenation responses to exercise are presented in Table 3 and Figs 4, 5, 6 and 7. Differences were *unclear* between INSP and CTRL for TSI_RM_ (1.0%, 90% CL ±7.5%), but MATCH was *very likely* lower than INSP (22.1% ±12.5%). Average HHb_RM_ was *likely* greater during INSP compared to CTRL (9.0%, 90% CL ±7.5%). Conversely, HHb_RM_ was lower during MATCH compared to INSP (−19.6%, 90% CL ±6.0%).

**Fig 4.**
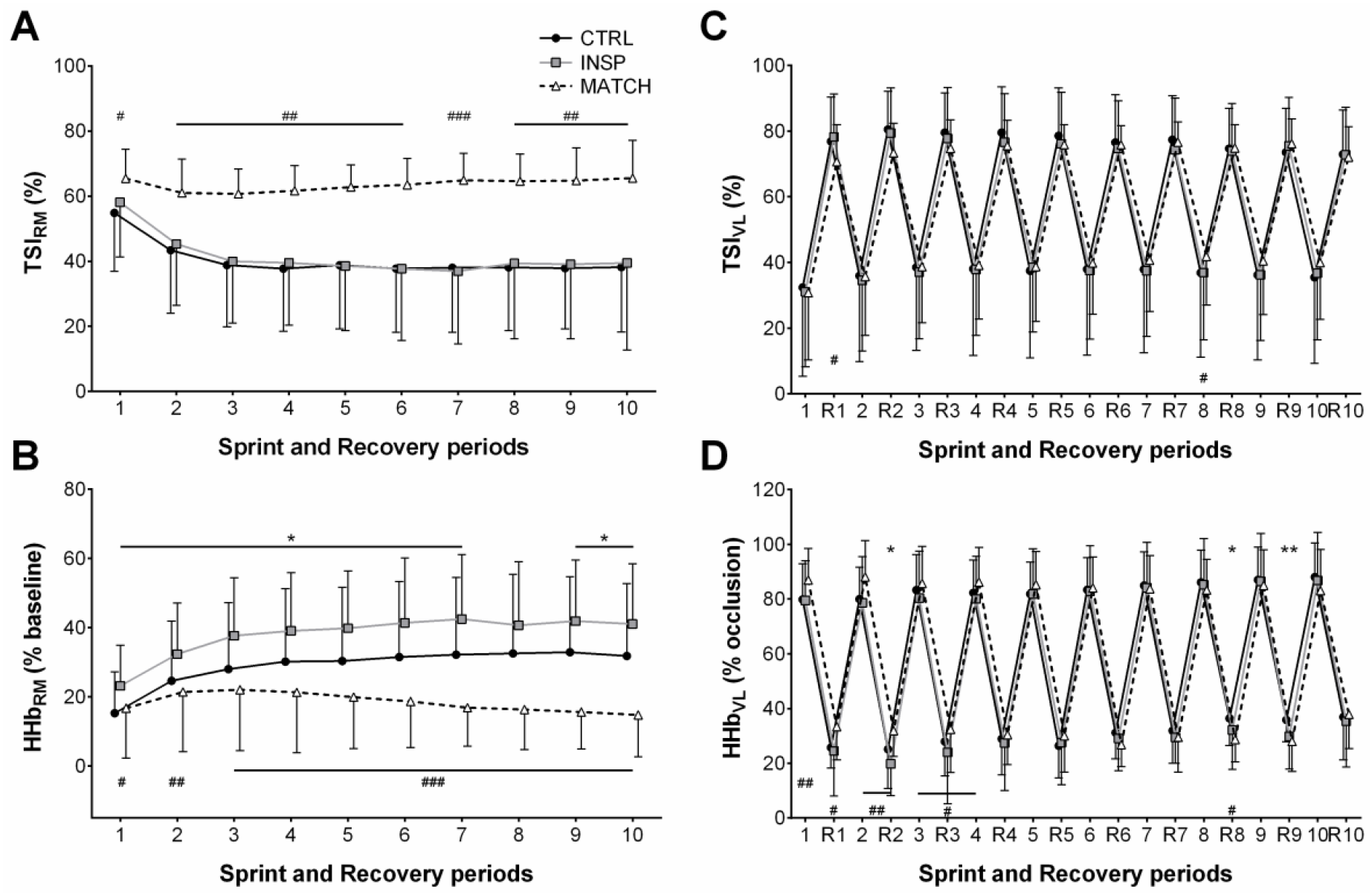
Respiratory and locomotor muscle NIRS responses to repeated-sprints during the control (CTRL) inspiratory loading (INSP) and work matched (MATCH) experimental sessions. (A) Respiratory muscle tissue saturation index (TSI_RM_) (n = 8). (B) Respiratory muscle deoxy-haemoglobin (HHb_RM_). (C) Vastus lateralis tissue saturation index (TSI_VL_). (D) Vastus lateralis deoxy-haemoglobin (HHb_VL_). Values are presented as mean ± SD. The symbols represent comparisons between INSP and Control (*), INSP and MATCH (#). The number of symbols; one, two and three denote likely, very likely and almost certainly respectively, that the chance of the true effect exceeds a small (−0.2-0.2) effect size. Vales are presented as mean ± SD.

**Fig 5:**
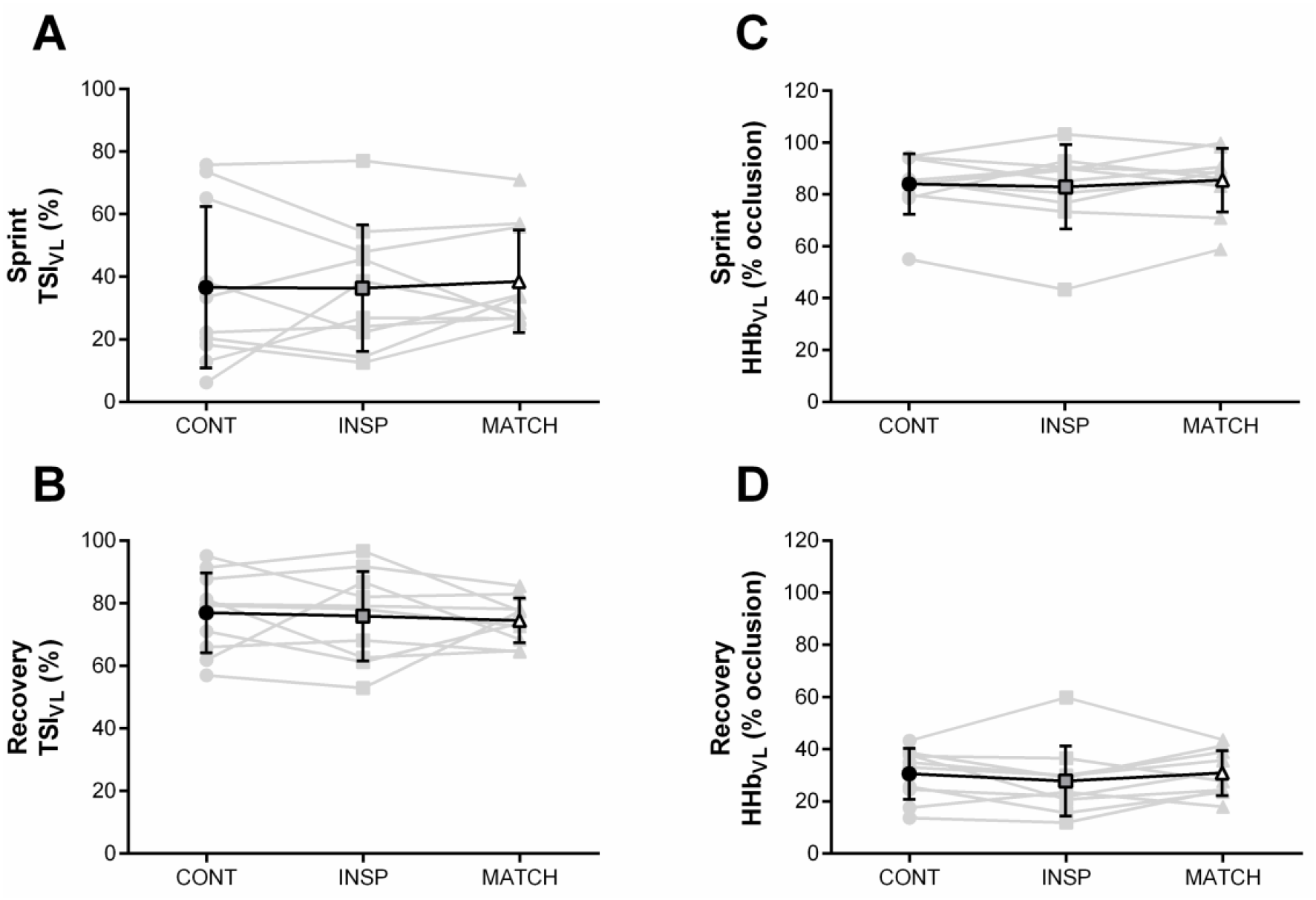
Mean locomotor muscle NIRS responses to repeated-sprints during the control (CTRL) inspiratory loading (INSP) and work matched (MATCH) experimental sessions. (A) Sprint vastus lateralis tissue saturation index (Sprint TSI_VL_). (B) Recovery vastus lateralis tissue saturation index (Recovery TSI_VL_). (C) Sprint vastus lateralis deoxy-haemoglobin (Sprint HHb_VL_). (D) Recovery vastus lateralis deoxy-haemoglobin (Recovery HHb_VL_). Individual responses are represented by the grey lines and mean responses are represented by black lines. The symbols represent comparisons between INSP and Control (□), INSP and MATCH (#). The number of symbols; one, two and three denote likely, very likely and almost certainly respectively, that the chance of the true effect exceeds a small (−0.2-0.2) effect size. Vales are presented as mean ± SD.

**Fig 6:**
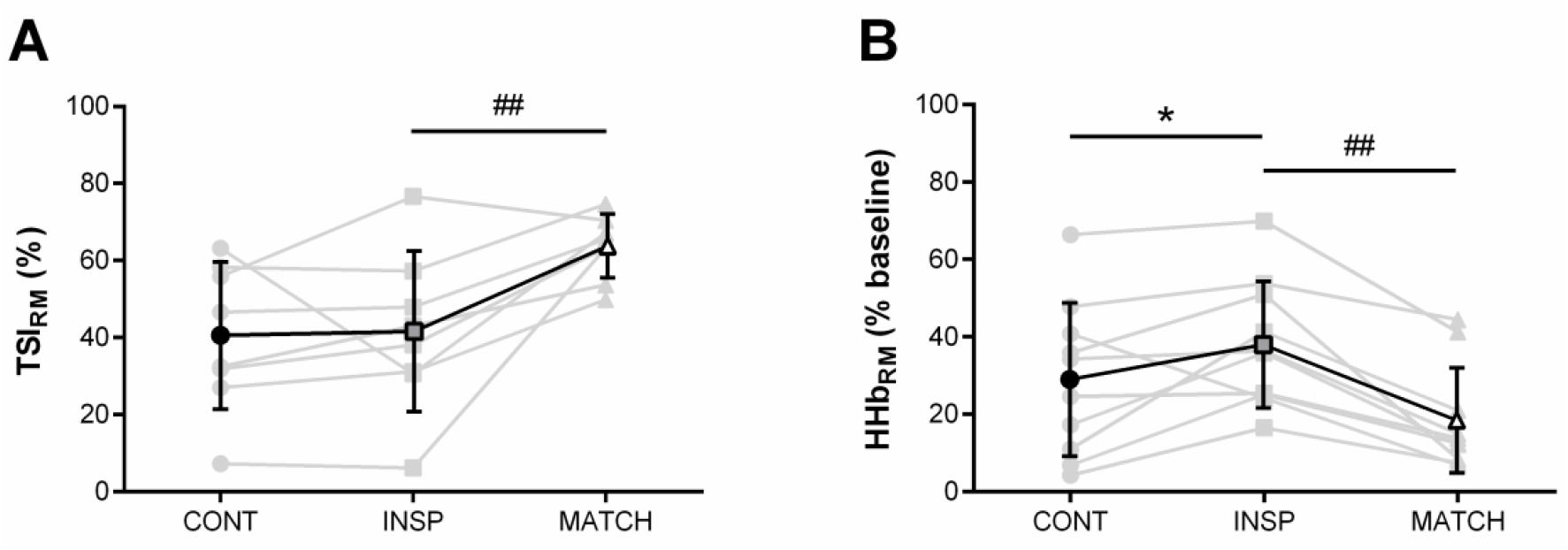
Mean respiratory muscle NIRS responses to repeated-sprints during the control (CTRL) inspiratory loading (INSP) and work matched (MATCH) experimental sessions. (A) Respiratory muscle tissue saturation index (TSI_RM_) (n = 8). (B) Respiratory muscle deoxy-haemoglobin (HHb_RM_). Individual responses are represented by the grey lines and mean responses are represented by black lines. The symbols represent comparisons between INSP and Control (□), INSP and MATCH (#). The number of symbols; one, two and three denote likely, very likely and almost certainly respectively, that the chance of the true effect exceeds a small (0.2-0.2) effect size. Vales are presented as mean ± SD.

**Fig 7:**
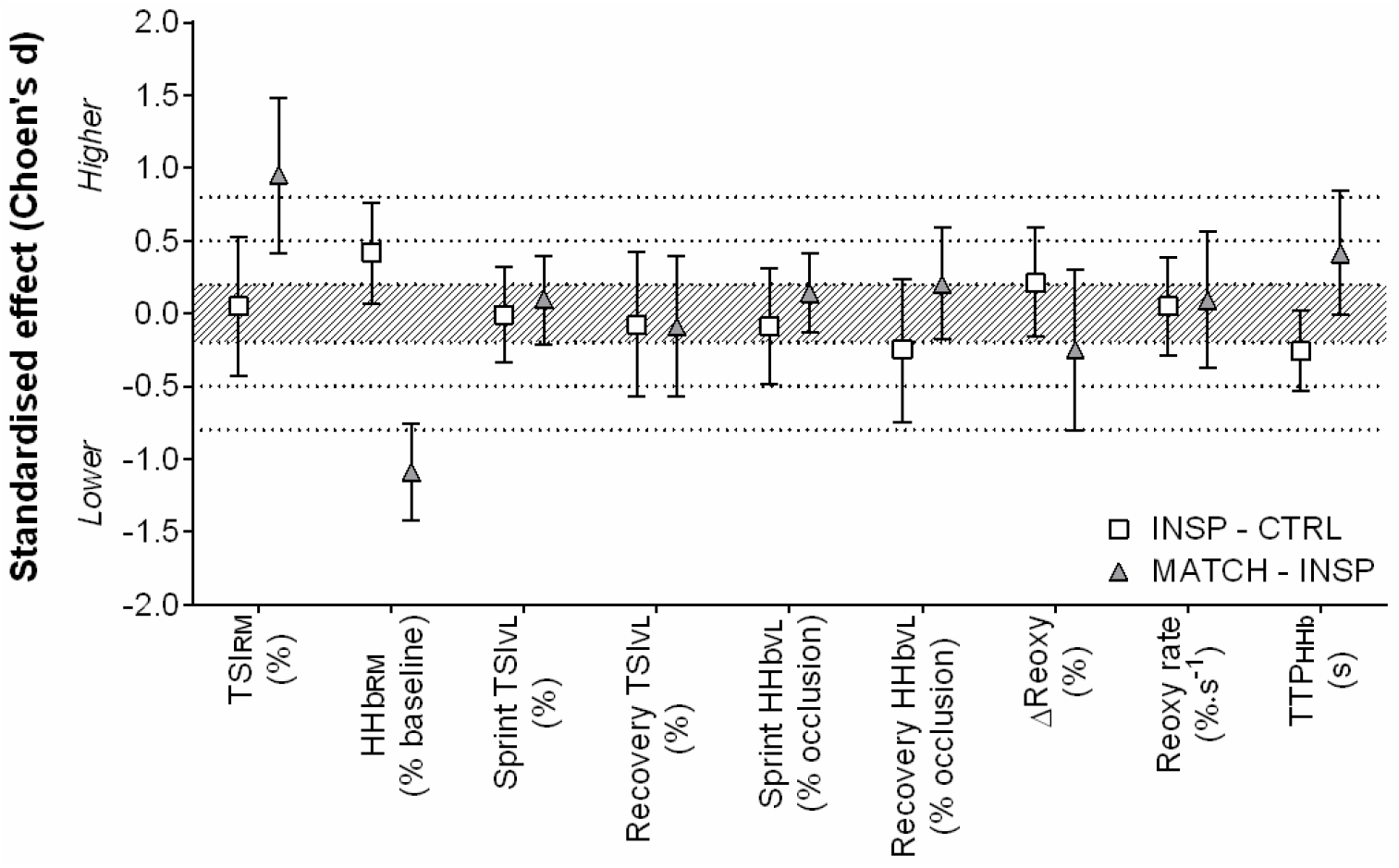
Standardised effects (Cohen’s d) with 90% confidence intervals for NIRS variables comparing inspiratory loading (INSP) to control (CTRL), and work matched exercise (MATCH) to INSP. Shaded area indicates a trivial effect, and dotted lines thresholds for small, moderate and large effects. Abbreviations are: respiratory muscle tissue saturation index, TSI_RM_; respiratory muscle deoxy-haemoglobin, HHb_RM_; vastus lateralis tissue saturation index, TSI_VL_; vastus lateralis deoxy-haemoglobin, HHb_VL_; vastus laterals reoxygenation, ΔReoxy; vastus laterals reoxygenation rate, Reoxy rate; time to peak deoxy-haemoglobin, TTP_HHb_.

**Table 3:** Near-infrared spectroscopy responses to repeated-sprint exercise during control (CTRL), inspiratory loading (INSP), and work match (MATCH) conditions.

| Variable | CTRL | INSP | MATCH |
| --- | --- | --- | --- |
| TTP<sub>HHb</sub> (s) | 13.0 ± 3.3 | 12.3 ± 3.4 | 13.8 ± 3.7 # |
| ΔReoxy (%) | 49.94 ± 20.30 | 54.66 ± 19.24 | 49.45 ± 21.08 |
| Reox rate (%·s<sup>-1</sup>) | 2.16 ± 0.78 | 2.20 ± 0.75 | 2.27 ± 0.79 |

Vales are reported as mean ± SD. Abbreviations are: TTP<sub>HHb</sub>; reoxygenation, ΔReoxy; reoxygenation rate, Reoxy rate. The symbols represent comparisons between INSP and CTRL (\*), INSP and MATCH (#). The number of symbols; one, two and three denote likely, very likely and almost certainly respectively, that the chance of the true effect exceeds a small (-0.2-0.2) effect size. Values are reported as mean ± SD.

Differences of sprint TSI_VL_ (−0.3% ±9.5%) and recovery TSI_VL_ (−1.1% ±7.0%) were *unclear* between INSP and CTRL. Similarly, the differences between INSP and CTRL for both sprint HHb_VL_ (−1.1% ±5.1) and recovery HHb_VL_ (−2.7% ±5.4%) were *unclear*. There was no meaningful difference between INSP and CTRL for TTP_HHb_ (−5.8% ±6.0%) and ΔReoxy (4.7% ±8.3). Additionally, there was an *unclear* difference in Reoxy rate (0.0% ±0.3%).

In MATCH exercise, differences in TSI_VL_ were unclear compared to INSP for both sprint (2.2% ±6.7%), and recovery (−1.4% ±7.6) phases. Additionally, there was no meaningful difference in sprint HHb_VL_ (−1.1% ±5.1%), and an *unclear* difference for recovery HHb_VL_ (3.0% ±5.7%). The TTP_HHb_ was greater during MATCH than INSP (12.0% ±14.4%). There were also *unclear* differences in ΔReoxy (−5.2% ±11.5%), and Reoxy rate (0.1% ±0.4%).

## Discussion

This study examined the respiratory and vastus lateralis muscle oxygenation trends during repeated-sprint exercise with heightened respiratory muscle work. The addition of inspiratory loading increased mouth pressure and respiratory muscle O_2_ deoxygenation.

However, these altered responses had no meaningful impact on blood arterial O_2_ saturation and tissue oxygenation trends within the vastus lateralis muscle. We interpret these findings to suggest that during maximal intermittent work, O_2_ delivery to the respiratory and locomotor muscles can be maintained.

### Work of breathing and respiratory muscle oxygenation

Hyperpnoea during high-intensity exercise requires a considerable portion of whole-body 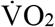 to support the metabolic demands of the respiratory muscles (16), and is increased when an inspiratory load is added (19). In the present study, pulmonary 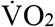 was elevated by 4-5% during both the sprint and recovery phases of the repeated-sprint protocol when an inspiratory load was added. This occurred even though there was no meaningful difference in total locomotor work completed during the INSP and CTRL experimental sessions.

When subjects exercised with the inspiratory load, 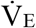 was 19.6% lower compared to CTRL, achieved via a reduction in *f*_R_. Regardless of these changes, S_P_O_2_ was not dissimilar between the exercise conditions. Either consciously or subconsciously choosing a lower 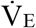, subjects may have counteracted or at least attenuated the expected work of breathing with INSP. When the change in *f*_R_ was accounted for, respiratory muscle force developments (∫P_m_ × *f*_R_) was substantially higher during INSP compared to CTRL (734%). Higher ∫P_m_ × *f*_R_ in conjunction with the elevated HHb_RM_ in the current study, suggests that O_2_ utilisation by the respiratory muscles was increased with inspiratory loading, as shown previously (20, 24). Furthermore, there were *unclear* differences in TSI_RM_ suggesting that [O_2_H_b_] was higher to match the demands for O_2_ delivery of the respiratory muscles. Previous studies have shown similar changes in HHbRM in response to inspiratory loading (24) and resistive breathing (25). The present data further suggests maintenance of respiratory muscle oxygenation with inspiratory loading which contrasts with others who reported similar values for f ∫P_m_ × *f*_R_ (33). However, preserved respiratory muscle oxygenation may have negative consequences for exercise tolerance if blood flow is redistributed away from the active limbs to meet the metabolic demands of breathing.

During continuous high-intensity exercise when the work of breathing is high, there can be an increase in vascular resistance and a reduction in limb perfusion (18, 19, 34). The accumulation of metabolites in the respiratory muscles stimulates group IV afferent discharge in these muscles (35), leading to sympathetically mediated efferent discharge and vasoconstriction in the locomotor muscles (18, 36). Despite the clear increase in respiratory muscle deoxygenation shown in the present study as a result of inspiratory loading, there was no clear negative effect on vastus lateralis oxygenation. The intermittent nature of repeated-sprint exercise may allow sufficient recovery time to prevent the accumulation of fatigue inducing metabolites and recover O_2_ debt in the respiratory muscles.

### Locomotor muscle oxygenation

Inspiratory loading had no discernible effects on sprint HHb_VL_ or TSI_VL_ despite a considerable increase in the work of breathing and respiratory muscle O_2_ utilisation. Vastus lateralis deoxygenation rapidly increases at sprint onset, and then plateaus with sprint repetitions (1, 11, 12). This suggests that a maximal level of O_2_ extraction in the locomotor muscles is achieved in “normal” exercise conditions (37). However, a higher secondary ceiling point to vastus lateralis deoxygenation has been observed when repeated-sprint exercise has been performed in simulated altitude (normobaric hypoxia) (13). Elevated muscle deoxygenation during maximal exercise may be compensatory for reduced muscle O_2_ availability (38). If vastus lateralis O_2_ availability had been impacted in the present study by an elevated work of breathing, sprint HHb_VL_ would have increased during INSP compared to CTRL. Nevertheless, vastus lateralis deoxygenation during the sprint phase per se may play a limited role in prolonged repeated-sprint performance. Muscle O_2_ availability during the recovery phase appears to be far more influential in maintaining performance as sprints are repeated (13). The capacity to reoxygenate muscle tissues between sprints is highly sensitive to O_2_ availability and underpins metabolic recovery between sprint bouts (11, 13, 15). Even with the addition of an inspiratory load in this study, which increased respiratory muscle O_2_ utilisation, vastus lateralis O_2_ delivery was maintained. It therefore appears that the cardiovascular system can support the metabolic O_2_ demands of both the respiratory and locomotor muscles during repeated-sprint exercise.

It is likely that locomotor muscle oxygenation will only be compromised when cardiac output can no longer increase to meet the O_2_ demands of both the respiratory and locomotor muscles simultaneously. It has been demonstrated that while exercising at near-maximal work rates, that there is no accompanying increase in cardiac output and 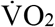 with inspiratory loading, and as a result limb blood flow is compromised (18, 19). This presumably occurs when the prescribed exercise intensity is sufficient to elicit sustained 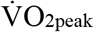, and therefore, no further increase can occur. While in the present study there was a 4-5% increase in 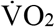 throughout the repeated-sprint protocol with inspiratory loading, which is similar to others who have shown an increase in 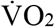 during submaximal exercise (20). These data demonstrate that during repeated-sprint exercise, there is available capacity for 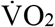 to increase to meet the heightened metabolic demands of inspiratory loading, which may be a crucial factor in maintaining O_2_ supply to the locomotor muscles. Secondly, the intermittent nature of repeated-sprint exercise will minimise the development of diaphragm fatigue and the activation of the respiratory muscle metaboreflex which is promoted during sustained high-intensity exercise (17). As shown in Fig 3, pulmonary 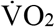 fluctuated between 90% and 70% of 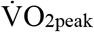 during sprint and recovery phases, respectively, during the control condition. Moreover, the addition of an inspiratory load increase 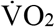 by ≈5% in both the sprint and recovery phases. The fluctuation in metabolic demands exemplified by the undulating 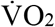 likely minimises the opportunity for substantial competition of available cardiac output to develop, and the impairment of tissue reoxygenation.

Hyperventilation was present in both CTRL, and to a lesser degree, INSP experimental sessions, which may have had a protective effect on limb O_2_ delivery. Hyperventilation is associated with an increase in alveolar ventilation disproportionate to 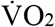 (pressure of alveolar O_2_ increases), and P_A_CO_2_ (pressure of alveolar CO_2_ decreases) (39). This is a potential mechanism associated with high-intensity exercise which can constrain a fall in arterial O_2_ and pH (39, 40). Despite a reduction of 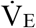 in a state of heightened O_2_ demand during the INSP session, S_P_O_2_ was maintained. In studies where vastus lateralis tissue oxygenation was impaired during exercise with resistive breathing and inspiratory loading, there was moderate exercise induced arterial hypoxemia (24, 25, 41). Exercise-induced arterial hypoxemia is a known limiting factor of exercise (41), and preventing it with supplemental O_2_ can attenuate peripheral muscle fatigue (42). Though exercise-induced arterial hypoxemia has been observed during longer repeated-sprint exercise (12), there was no evidence of its occurrence in the present study. Meaningful changes in repeated-sprint oxygenation trends from an elevated work of breathing may only occur if a moderate level exercise-induced arterial hypoxemia (88-93%) is also present. Some caution should be expressed when interpreting fingertip S_P_O_2_ which we and others have used (25), since it is prone to artefact and data loss from movement and tight gripping of the cycle ergometer handle bars (43). Monitoring S_P_O_2_ from the earlobe (24, 44) would have overcome this limitation and increased the accuracy of our measurements.

The level of inspiratory loading may have also had an influence on the outcomes in this study. In previous work, inspiratory loading was achieved by reducing the inspiratory aperture to 10 mm and 8 mm (24). Changes in [HHb] of the exercise limb were only detected with the smaller opening. Similarly, when resistive breathing has been used, the most noticeable changes in tissue oxygenation trends occurred when the aperture was reduced to 4.5 mm (25). One may argue that the inspiratory work in the present study was too low to induce a respiratory muscle metaboreflex, however peak P_m_ and ∫P_m_ × *f*_R_ was similar to the previous work using an 8-mm aperture (24, 33). Additionally, 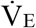 was considerably higher than in previous work (24, 25), which would have contributed to a heightened work of breathing.

### Work-matched exercise

To our knowledge, this is the first-time repeated work matched bouts of exercise have been used to examine the demands of repeated-sprint exercise under altered metabolic conditions. Despite a similar degree of vastus lateralis tissue deoxygenation during the work-matched sprints, the physiological load placed on the cardiovascular system was considerably lower. This is evidenced by the consistently lower 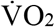, probably due to markedly lower respiratory muscle O_2_ utilisation. But more importantly, 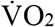 was heavily influenced by how exercise was prescribed. Matching total work was achieved by replicating mean power output for each sprint, and therefore was lacking maximal acceleration and power production typically associated with sprint exercise. For example, there was a clear difference in peak power output between INSP (1097 ± 148 W) and MATCH (773 ± 122 W) exercise conditions. In a typical sprint, power output peeks within 1-2 s and progressively declines as the sprint continues (23, 45). When sprints are repeated and separated by incomplete recovery periods, both peak and mean power output gradually decline as fatigue accumulates (1–4, 44). By matching for mean power output and eliminating the opptunity for subjects to “maximally” exert themselves, the reliance on intramuscular ATP and PCr hydrolysis would have been reduced (2, 46), and metabolic perturbations associated with maximal exercise minimised (2, 5, 46). Despite a substantial decrease in 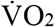 and the O_2_ cost associated with the work of breathing, there was no substantial difference in ΔReoxy nor Reoxy rate. This implies that tissue reoxygenation was maximal in all exercise conditions since the now “available” cardiac output was not being utilised to reoxygenate the locomotor muscles (18). It appears that there exists some degree of reserve in the cardiovascular system that is called upon to maintain O_2_ delivery to both respiratory and locomotor muscles when the work of breathing is high. Therefore, the O_2_ cost of breathing in repeated-sprint cycling is unlikely to have a meaningful negative impact on locomotor O_2_ transport.

## Conclusions

A crucial factor of repeated-sprint performance is the reoxygenation capacity between sprint bouts (11, 13, 14). We further tested this mechanism by increasing the work of breathing, which is known to negatively influence limb blood flow and O_2_ delivery at least in endurance exercise (16–18, 34). The present data demonstrate that the addition of inspiratory loading did not impair oxygenation to the vastus lateralis. When maximal exercise is interspersed with short rest periods, the cardiovascular system appears to maintain O_2_ delivery, to both the locomotor and respiratory muscles in a state of heightened metabolic demand.

